# Relaxed risk of predation drives parallel evolution of stickleback behaviour

**DOI:** 10.1101/2022.01.05.475046

**Authors:** Antoine Fraimout, Elisa Päiviö, Juha Merilä

## Abstract

The occurrence of similar phenotypes in multiple independent populations (*viz*. parallel evolution) is a testimony of evolution by natural selection. Parallel evolution implies that populations share a common phenotypic response to a common selection pressure associated with habitat similarity. Examples of parallel evolution at the genetic and phenotypic levels are fairly common, but the driving selective agents often remain elusive. Similarly, the role of phenotypic plasticity in facilitating early stages of parallel evolution is unclear. We investigated whether the relaxation of predation pressure associated with the colonization of freshwater ponds by nine-spined sticklebacks (*Pungitius pungitius*) likely explains the divergence in complex behaviours between marine and pond populations, and whether this divergence is parallel. Using laboratory-raised individuals exposed to different levels of perceived predation risk, we calculated vectors of phenotypic divergence for four behavioural traits between habitats and predation risk treatments. We found a significant correlation between the directions of evolutionary divergence and phenotypic plasticity, suggesting that habitat divergence in behaviour is aligned with the response to relaxation of predation pressure. Finally, we show that this alignment is found across multiple pairs of populations, and that the relaxation of predation pressure has likely driven parallel evolution of behaviour in this species.

## INTRODUCTION

Similar environments may impose similar selection pressures on newly colonizing populations, leading to recurrent phenotypes in multiple habitats (Bailey *et al*. 2015). The evolution of similar phenotypes between lineages – convergent evolution (Rosenblum *et al*. 2014) – has long been attributed to natural selection, as only such a deterministic process is expected to result in the occurrence of the same traits in similar environments (Rundle *et al*. 2000, Schluter *et al*. 2004). Recent studies of repeated evolution in the wild have greatly advanced our understanding of the population-specific factors influencing the likelihood of parallel evolution (Stern & Lee 2020, Fang *et al*. 2021, Kingman *et al*. 2021) and the genetic underpinnings of convergent phenotypic adaptation (Xie *et al*. 2019, Kemppainen *et al*. 2021). Nonetheless, these detailed studies of the genetic mechanisms involved in the response to selection, often elude identifying the actual selective agents behind the observed responses. Yet, the main premise of repeated evolution is that the lineages evolving in parallel should do so in response to a common selection pressure and therefore, identifying the environmental factors driving these responses is central to our understanding of parallel evolution.

Predation is a ubiquitous feature of ecosystems and a driving force of the evolution of species interactions (Abrams 2000). Because of its direct influence on fitness, predation is also a strong selective agent behind the evolution of morphological (Eklöv *et al*. 2006), physiological (Rödl *et al*. 2007) and behavioural traits (Lapiedra *et al*. 2018) in prey species. While predation can shape the distribution of phenotypes in prey communities, relaxation of predation pressure – *e*.*g*., following the colonization of a predator-free habitat – has been suggested to favour certain traits and lead to the evolution of novel phenotypes (Bliard *et al*. 2020). In either case, the presence or absence of predators in the environment is expected to play a central role in adaptive evolution, and generate long-term divergence stemming from different levels of predation (Nosil 2004, Nosil & Crespi 2006). Changes in the predation regime of an environment can also induce short-term individual responses through phenotypic plasticity (see West-Eberhard 2003 for definition and Benard 2004 for review). For instance, organisms may adjust their behaviour when predation risk is high to either increase their probability of survival (Wen & Ueno 2021), or the survival of their offspring (Peluc *et al*. 2008). Consequently, individual variation in the magnitude and direction of plasticity in a population provides an additional source of phenotypic variation for selection to act on (Abbey-Lee & Dingemanse 2019), and it has been hypothesized that plasticity can sometimes ‘take the lead’ in early stages of adaptive evolution (Scoville & Pfrender 2010, Levis & Pfennig, 2016, 2020). Empirical evidence for the role of phenotypic plasticity in repeated evolution of complex traits is still relatively scarce, yet its putative part in paving the way of adaptive evolution holds an important place in the Extended Evolutionary Synthesis (Futuyma 2017).

Here, we investigated the effects of perceived predation risk on the expression of behavioural traits in two types of locally adapted populations of the nine-spined stickleback (*Pungitius pungitius*). The nine-spined stickleback is a teleost fish distributed across the northern parts of Eurasia and North America. An ecological peculiarity of this species is that it naturally occurs in both marine and freshwater habitats. Marine ancestral populations of *P. pungitius* have colonized multiple freshwater habitats following the last glaciation ca. 11,000 years ago (Feng *et al*. 2021) and *P. pungitius* are now found in isolated ponds throughout Northern Europe (Teacher *et al*. 2011). Whereas marine populations of *P. pungitius* co-occur with a diverse community of piscine predators, freshwater pond populations have evolved in a virtually predator-free environment where they are often the sole fish species (Herczeg *et al*. 2010). As a result, it has been hypothesized that pond populations have evolved remarkable phenotypes in response to this relaxation of predation pressure, including gigantism (Herczeg *et al*. 2009a) and bold aggressive behaviours (Herczeg *et al*. 2009b). Empirical evidence demonstrated that among-habitat differences in behaviour are genetically based and have resulted from divergent selection acting on several behavioural traits (Karhunen *et al*. 2014). Despite this evidence, whether predation is the likely factor driving behavioural divergence among habitats, and whether such divergence has repeatedly occurred in parallel, is yet to be tested experimentally.

We hypothesized that the relaxation of predation pressure associated with the colonization of predator-free habitats has driven the evolution of behaviour in pond populations of *P. pungitius*.

Furthermore, we test the complementary hypothesis that the between-habitat divergence in behaviour may have resulted from the expression of advantageous plastic phenotypes in response to the relaxation of predation pressure. To this end we used an experimental test of behavioural response to predation exposure in pond and marine nine-spined sticklebacks, and addressed the following questions: i) Did behaviour evolve in parallel among freshwater *P. puntitius* populations? To answer this question, we verified the expectation that parallel evolution of behaviour should be reflected by an alignment between the phenotypic vectors of divergence from a marine ancestor, between multiple pond populations. ii) Is the relaxation of predation pressure likely to be the selective agent underlying the divergence between marine and pond sticklebacks? For this, we tested the theoretical prediction (Lind *et al*. 2015, Radersma *et al*. 2020) that the vector of phenotypic plasticity stemming from our predation exposure treatment should be aligned with the vector of phenotypic divergence between habitats.

## MATERIALS AND METHODS

### Sampling

Adult *P. pungitius* were sampled during breeding season (May – June 2018) at eight different locations in Finland and Sweden corresponding to four coastal marine and four freshwater pond habitats (Table. S1). Pond populations were sampled using minnow traps placed in ca. 50 cm depth and marine populations were sampled from shallow (ca. 1m depth) waters using beach-seine nets. Sampled fish were checked visually to ensure sexual maturity (*i*.*e*., black abdomen in males and rounded bellies in gravid females, *e*.*g*., McLennan, 1996) and subsequently transported to the aquaculture facilities of the University of Helsinki. Wild-caught individuals from each population were housed separately in 1m^3^ plastic aquaria with flow-through water system and fed *ad libitum* with frozen chironomid larvae twice a day.

### Common garden experiments

In order to control for environmental variance and to measure genetically-based phenotypic variation among individuals, we set up a common-garden rearing design in the laboratory: for each population, 5 to 10 full-sib families were produced (*n* = 65; Table S1) by artificial crossing of random pairs of wild-caught individuals. We followed the standard *in vitro* fertilization techniques and egg husbandry protocols for stickleback crossing (Arnott and Barber, 2000) and obtained eggs from gravid females by gently squeezing their abdomens over a petri dish. Males were over-anesthetized using tricaine methanesulfonate (MS-222) in order to dissect their testes, which were subsequently minced in the petri dish containing the eggs. Eggs and sperm were gently mixed using a plastic pipette to ensure fertilization, and kept in water until hatching. Water in the petri dishes was changed twice a day and clutches were visually checked for signs of fungal infections or death, and accordingly removed. At the onset of hatching and for a four weeks period, each clutch was split in two replicate 11 × 10 cm plastic boxes. Following yolk resorption, fry was fed *ad libitum* with live brine shrimp (*Artemia sp. nauplii*). All replicated families were transferred to Allentown Zebrafish Rack Systems (hereafter rack; Aquaneering Inc., San Diego, USA). Racks had a closed water circulation system, with multi-level filtering including physical, chemical, biological and UV filters. All fish were reared in racks under constant temperature and light conditions (15°C; 12:12 LD) for a period of ca. 1 year (mean age: 316.4 days) until the start of the behavioural experiment. We ensured that all fish did not show signs of sexual maturity which could affect the expression of behaviours. Before starting the experiments, all families were transferred to holding tanks where they were kept in constant temperature and light conditions (15°C; 12:12 LD) throughout the experimental periods. Replicates of the same family were housed in separate tanks in order to account for common environment variance.

### Experimental setup

Two identical experimental aquaria with independent flow-through water systems were built for the experiments (*Supplementary methods*; Fig. S1). Each aquarium was divided transversely in two sections by a transparent plastic plate separating the behavioural arena and the holding arena. The behavioural arena corresponded to the half of the tank where the focal fish were placed and scored for behaviours, while the holding arena corresponded to the half where the predators were introduced (predation treatment) or left empty (control treatment; see below). In order to investigate the effect of predation risk on stickleback behaviour, behavioural tests were conducted in the presence and absence of predators. One of the experimental aquaria was assigned to predation treatment and one to control treatment. In the predation treatment, a pair of wild-caught perch (*Perca fluviatilis)*, a natural predator of marine *P. pungitius* (Nelson & Bonsdorff 1990), were placed on the holding arena of the experimental aquarium.

### Behavioural measurements

We measured ecologically relevant behaviours classified into two categories: exploration (an individual’s propensity to explore a novel environment), and risk-taking during foraging (an individual’s tendency to take risks to obtain food). All behavioural measurements were performed with one fish at a time and fish were starved for 24 hours prior to the experiments. Each trial started by transferring the focal fish from the holding tank into the behavioural arena of the experimental tank and running the exploration test followed by the risk-taking test (see also *Supplementary methods* for details).

The focal fish was caught from its holding tank with a hand net and introduced into the cylinder in the experimental tank (Fig. S1). The fish was left to acclimatise inside the cylinder for three minutes. After this acclimation time, the door of the cylinder was opened allowing the fish to leave the cylinder to explore the experimental tank. Two measurements were recorded: the latency until the head of the fish came out of the cylinder, and the latency until the full body of the fish came out of the cylinder.

Following the exploration test, the cylinder was removed, and the fish was left to acclimatize for three minutes in the behavioural arena. After the acclimation period, chironomid larvae (a familiar food) were pipetted into the open area of the tank in a straight diagonal line from the edge of the refuge to the opposite corner of the tank (see Fig. S1). With this kind of food administration, the more the fish ate, the further it had to move from the refuge, so that the “risk” experienced by the fish (swimming further into the open area and closer to the predator) was proportional to the “reward” (number of food items). Three measurements were recorded: the time spent in the open area (whole body outside the refuge area when viewed from above) in the five minutes following the addition of the first food item; the latency to initiate feeding after the addition of the first food item; and the total number of feeding events measured as the number of successful attacks on the food.

All time variables (latencies) were measured in seconds and each trial was terminated if the fish did not express the behaviour after 5 minutes, so that the maximum value for these measurements was 300 seconds. At the end of the experiment, a total of 422 individuals were phenotyped across 65 families and eight populations for the four following traits: emergence time (the arithmetic mean of time-to-head-out and time-to-body-out variables, see *Supplementary methods*), open time (time spent in the open area), feeding (the number of feeding bouts) and risk-taking (the latency to first feeding).

### Statistical tests of phenotypic differentiation

We first investigated behavioural variation between populations, habitats and the effect of perceived predation using statistical models. Our data consisted of three right-censored (*i*.*e*., truncated) time-to-event variables. This type of data is not suitable for classical linear regression approaches (*i*.*e*., linear mixed-or generalized linear models; Edelaar *et al*. 2012) and we thus followed multiple statistical frameworks to verify the robustness of our results (see *Data analysis of right-censored data* in *Supplementary methods*). We here present the main analysis applied to these variables. For the right-censored time-to-event variables (*i*.*e*., emergence time, open time, risk-taking), we fitted censored regressions using the *censReg* R package (v.0.5-32, Henningsen 2017). Main fixed effects of interest included habitat of origin and treatment (predation or control) and their interaction, and setting the right limit for censoring at 300 (the maximum time value in seconds in our experiment).

Count data (*i*.*e*., feeding variable) were analysed with a generalized linear model (GLM) using the *glm* function in the *lme4* R package (v.1.1-27, Bates *et al*. 2015) with habitat of origin and treatment and their interaction as fixed effects. To account for the possible effects of body size and age variation in our data, we fitted all the above models including an age-corrected body size covariate, computed from the residuals of a linear regression of body size on age. Temporal block of measurements (see *Supplementary methods*) was set as fixed effect in all models.

### Phenotypic vector analysis

We investigated parallelism in behavioural evolution by computing two types of phenotypic vectors: first, we estimated the evolutionary divergence vectors (Δ*z*_*D*_) corresponding to the phenotypic differences between marine and freshwater habitats. Specifically, we calculated the vectors of phenotypic change between each pond population from a hypothetical marine ancestral population. The ancestral marine population was estimated as the average behavioural phenotype from all the marine individuals measured in the presence of predators. We used these measurements as representative of a natural marine population experiencing predation pressure. Following the same logic, pond populations in the control treatment (no predation) were used as representative of natural freshwater populations experiencing no piscine predation. Vectors were calculated as the phenotypic difference between each pond population and the hypothetical ancestral population such that:

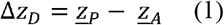

where *<u>z</u>_P_* corresponds to the mean phenotype of a pond population and *<u>z</u>_A_* to the mean phenotype in the ancestral marine population. Mean population phenotypes were extracted from separate model coefficients (censored regression or GLM, see above) using each behaviour trait as response variable and population of origin, treatment and their interaction as fixed effects. Age-corrected body sizes were used as covariates in all models as described above.

Second, we estimated the vectors of phenotypic plasticity (hereafter, plasticity vectors, *Δz_φ_*) as the phenotypic change induced by predation exposure. We were primarily interested in the plasticity vectors depicting the behavioural changes following the relaxation of predation pressure and thus, equivalent to the colonization of predator-free freshwater habitats by historical marine *P. pungitius* populations. To this end, we calculated the plasticity vectors as the phenotypic changes between the hypothetical ancestral population and each marine population measured in the control treatment as:

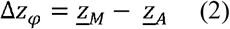

where *<u>z</u>_M_* is the mean trait value for the marine population measured in the absence of predators and *<u>z</u>_A_*, is the same as in eq. (1).

In order to test for the alignment between all pairs of divergence and plasticity vectors, we estimated the angle θ between any two pairs of vectors as:

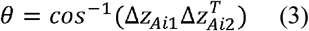

where each vector *Δz* corresponds to the normalized phenotypic vector of difference between the focal population *i* and the estimated marine ancestor *A*. Angles were calculated in degrees between all pairwise combinations of divergence and plasticity vectors. We assessed the statistical significance of all observed angles by comparing them to the angles calculated from 10,000 random vectors drawn from a normal distribution. Because we were interested in evaluating the evolution of complex behaviour in *P. pungitius*, each phenotypic vector described above was constructed from the multivariate behavioural traits’ dataset in each population and treatment. In other words, each vector of divergence or plasticity included the differences in means for all four behaviour traits measured, thus providing a multivariate measure of differentiation. In order to avoid scaling issues due to the differences between count data (*i*.*e*., feeding behaviour) and time-to-event data, raw measurements were transformed to z-scores using the *scale* function in R (v.4.1.1, R core team, 2021) prior to all phenotypic vector analyses.

We then followed the methodology of De Lisle & Bolnick (2021) to identify the dimensions of parallel change among divergence and plasticity vectors by analysing **C**, the matrix of correlation between replicated pairs of phenotypic vectors. We started this by constructing the matrix **X**, an *m* x *n* matrix with *m* rows containing each pairwise divergence and/or plasticity vector (*i*.*e*. each Δ*z*_*φ*_ and Δ*z*_*D*_) and *n* columns containing each behavioural trait (in our case 8 × 4). **C** was calculated as:

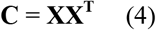

Eigenanalysis of **C** further allowed us to estimate whether one or more direction in the multivariate space (the eigenvectors) underlined a common parallel direction among our study populations, as well as the extent to which certain populations showed more parallelism among each other (see *Results* section) than others. All analyses were performed in R v.4.1.1 (R core team, 2021).

## RESULTS

### Phenotypic differentiation

There was a strong habitat differentiation in all behaviour variables and pond sticklebacks were consistently more explorative and took more risks during foraging than marine sticklebacks (Fig. 1A-D; Fig S2; Table S2). Overall, the predation treatment had stronger effects on foraging behaviours than exploration behaviour (Fig 1; Table S2-S4). Both pond and marine fish reduced the amount of feeding (Fig.1, Table S2) and took longer time to initiate feeding in the presence of predators (Fig. 1, Table S2-S4). Emergence time was not significantly affected by the presence of predators (Fig 1. Table S2-S4) but the predation treatment accentuated the habitat difference for this trait, with pond fish showing quicker emergence from refuge in the predation treatment (Fig. 1, Table S4). We found that our results were robust across different statistical methods (Table S2-S4) with the exception of open time: marine individuals were less likely to spend time in the open area in the presence of predators whereas the predation treatment did not lead to a significant decrease in open time in pond fish (Fig. 1, Table S2-3) but this result was not reflected by differences in survival curves using the Kaplan-Meier framework (Fig. S2, Table S4). We found limited statistical support for a significant interaction between the treatment and habitat effects in our models (only for open time using Box-Cox transformed data, Table S3), suggesting that both pond and marine fish had a similar plastic response to the exposure to predators. Finally, age-corrected body size only had a significant effect on the risk-taking behaviour with larger fish showing increased latency to first feeding (Table S2-S3).

**Figure 1.**
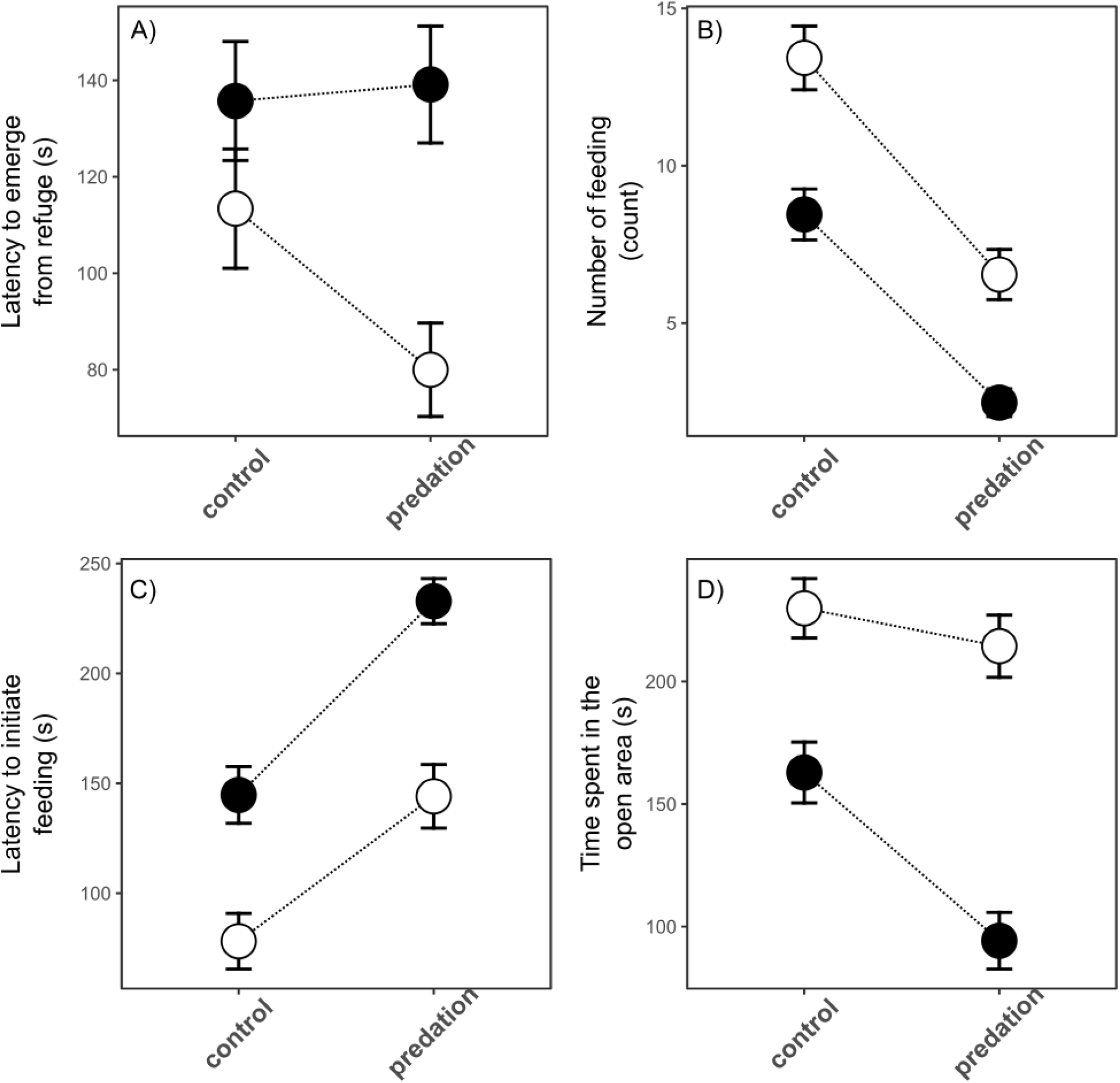
Behavioural variation between habitats and treatments. Mean values (circles) and standard errors (whiskered vertical bars) for the raw behaviour measurements are shown for marine (filled circles) and pond (open circles) fish in the control and predation treatments. A: Emergence time, the latency to emerge from a refuge (in seconds); B: Feeding, the number of feeding event (count); C: Risk-taking, the latency to initiate feeding (in seconds); D: Open time, the time spent in the open area (in seconds). Dashed lines represent the reaction norms for each habitat.

### Phenotypic vector analyses

Our phenotypic vector analyses allowed us to investigate three aspects of the evolution of complex behaviour in *P. pungitius* (Fig. 2, Table 1): i) the degree of parallelism between vectors of freshwater adaptation, indicated by the among-ponds comparisons of vectors (Table 1, green cells); ii) the degree of parallelism between the vectors of phenotypic plasticity, indicated by the among-marine comparisons of vectors (Table 1, blue cells) and iii) the correlation between the vectors of plasticity and evolutionary divergence, indicative of the effect of predation relaxation on the evolution of behaviour from marine to pond habitats (Table 1, red cells). We found that three out of the four pond populations shared a parallel direction of phenotypic divergence from the ancestral marine population, as evidenced by the small angles between their divergence vectors, which were found to be more similar than between random vectors (Table 1). The FIN-KRK population consistently showed evidence for non-parallelism with the other pond populations (Table 1, Fig. 1). We found that the plastic response to the relaxation of predation pressure was largely shared among marine populations.

**Table 1.** Angles between phenotypic vectors. The angle in degrees (above diagonal) between each pairwise vector comparison is shown along with their corresponding *p-values* (below diagonal) testing for significant differences between observed and random vectors (see *Methods*). Colour shading indicates the pairwise comparisons related to the test of parallel evolution among ponds (green), parallel phenotypic plasticity (blue) and the alignment between plasticity and divergence vectors (red, and see *Methods* for rationale). Bold values indicate statistical significance (*p* < 0.05) and italic values non-significance.

|  | FIN-PYO | FIN-RYT | FIN-KRK | SWE-BYN | FIN-TVA | FIN-POR | FIN-RAA | SWE-UME |
| --- | --- | --- | --- | --- | --- | --- | --- | --- |
| FIN-PYO |  | **7.144** | 77.850 | **15.310** | **23.694** | *35.151* | **9.694** | **23.438** |
| FIN-RYT | **<0.001** |  | 71.676 | **11.130** | **20.409** | *30.321* | **15.382** | **17.105** |
| FIN-KRK | *0.734* | *0.608* |  | *63.316* | *56.894* | *44.654* | *80.025* | *56.137* |
| SWE-BYN | **0.008** | **0.003** | *0.456* |  | **11.655** | **22.085** | **18.414** | **11.006** |
| FIN-TVA | **0.029** | **0.020** | *0.345* | **0.004** |  | **13.021** | **23.304** | **17.207** |
| FIN-POR | *0.093* | *0.059* | *0.178* | **0.021** | **0.005** |  | *36.020* | **21.079** |
| FIN-RAA | **0.003** | **0.010** | *0.781* | **0.015** | **0.029** | *0.101* |  | *28.883* |
| SWE-UME | **0.030** | **0.011** | *0.331* | **0.003** | **0.012** | **0.022** | *0.053* |  |

**Figure 2.**
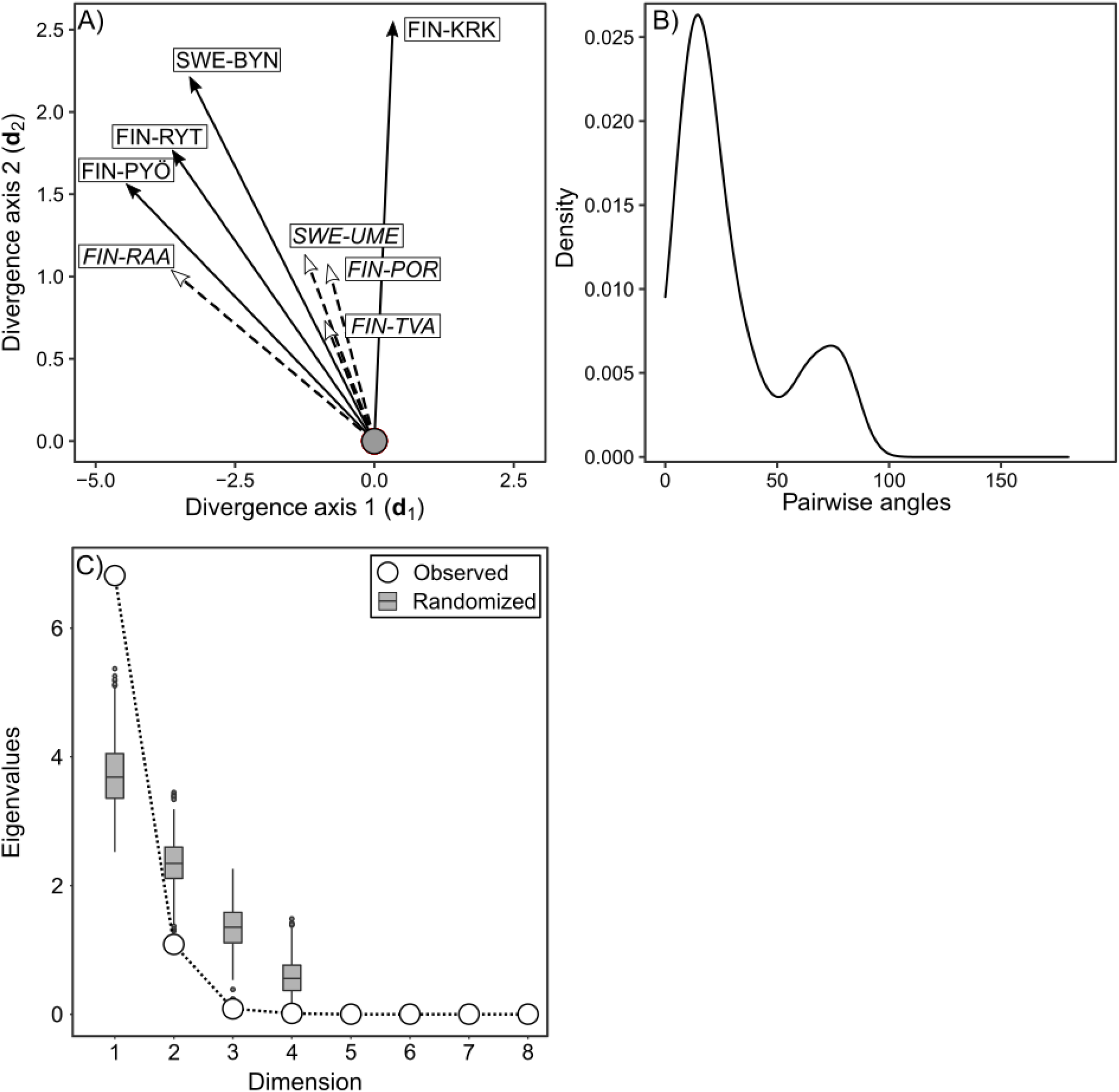
Results of the phenotypic vector analyses. A) Graphical representation of the phenotypic vectors. The vectors of divergence (filled solid arrows) and plasticity (open dashed arrows) from the ancestral marine population (gray filled circle) are projected in the multivariate divergence space where d1 and d2 represent the first and second main axis of the multivariate divergent covariance matrix. Population codes for pond and marine (italic) are indicated in black text. B) The distribution of observed vector angles in degree. C) Results of the multivariate test of parallelism. Eigenvalues from the decomposition of the **C** matrix calculated from the observed (open circles) and randomized (gray boxplots) data are shown. Boxplots represent the expected (randomized) eigenvalues calculated from sampling a Wishart distribution. Observed eigenvalues greater than expected ones indicate a single significant axis of parallelism among vectors.

Only one pair of populations (FIN-POR and FIN-RAA, Table 1) did not show evidence of parallelism between the vectors of phenotypic plasticity and another pair (FIN-RAA and SWE-UME, Table 1) had a small but marginally non-significant angle between vectors. Out of the 16 pairs of plasticity-divergence vectors, 10 showed significant parallelism, as indicated by the low angles between each pair of vectors (Table 1). The six non-significant parallel pairs of vectors all included the FIN-KRK and FIN-POR populations, indicating that the divergence of the FIN-KRK population from the marine ancestor did not follow the global direction of phenotypic plasticity and, conversely, that the plastic response of the FIN-POR population, did not align with the divergence vectors of all pond populations (Table 1). Overall, alignments between divergence and plasticity vectors indicate that the direction of behavioural change in the multivariate trait space induced by the relaxation of predation is similar to the direction of change observed in nature between marine and pond habitats.

Finally, we found that the directions of phenotypic changes stemming from the between-habitat divergence and the experimental relaxation of predation treatment were underlined by a single orthogonal dimension or parallelism, as evidenced by the first dimension of the **C** matrix decomposition (Fig. 2) showing greater eigenvalue than expected at random.

## DISCUSSION

Our common garden experiment shows that genetically-based differences in behaviour among pond and marine populations of *P. pungitius* have repeatedly evolved in parallel from marine ancestors. We found that our predation treatment generated a strong plastic response in most behavioural traits in both habitats and that this plastic response was aligned with the direction of evolutionary divergence. Below we discuss the implications of our results for the study of behavioural evolution in the wild.

The analyses of phenotypic vectors were based on a hypothetical marine ancestral population, corresponding to the average behavioural phenotype of contemporary Baltic Sea populations of *P. pungitius*. The detailed phylogeographic history of the nine-spined sticklebacks in Fennoscandia was recently resolved (Feng *et al*. 2021) and suggests that the Finnish pond and northernmost Baltic marine populations used in the current study most likely originated from ancestral populations in the White Sea rather than from the Baltic Sea (Teacher *et al*. 2011, Bruneaux *et al*. 2013, Feng *et al*. 2021). Nonetheless, Baltic *P. pungitius* are expected to be phenotypically similar (particularly regarding behaviour) to contemporary populations found in the White Sea (Herczeg *et al*. 2009a, Karhunen *et al*. 2014). More importantly, statistical modelling of behavioural phenotypes in relation to genetic coancestry revealed that the behaviour of contemporary marine populations of *P. pungitius* (Baltic and White Sea) is akin to the expected ancestral marine behaviour (see Fig. 3C, D in Karhunen *et al*. 2014). Our reconstruction of the ancestral population in the current analyses should thus be valid.

Pairwise comparisons of phenotypic vectors showed that the divergence of one freshwater population (FIN-KRK) deviated from that of other pond populations. Although we did not record the presence of other fish species at the time of sampling at this location, artificial introduction of potentially predatory trout (*Salmo trutta*) has been reported in this pond (Herczeg *et al*. 2010), and could explain the observed divergence in behaviour of this population. We also note that this population had the lowest sample size of our study and that estimates may be biased. Nevertheless, our multivariate test of parallelism identified a shared direction of phenotypic divergence among all pond populations, providing good evidence for the parallel evolution of behaviour associated with the colonization of freshwater habitat in this species. Moreover, this shared direction of parallelism also indicated that the direction of phenotypic plasticity generated by our control treatment (relaxation of predation pressure) is aligned with the direction of evolutionary divergence among habitats.

As for any other trait, evolution of phenotypic plasticity would require that the plastic response is genetically based and variable between individuals, and that this response would be advantageous in the environment where it is expressed (Ghalambor *et al*. 2007). Here, we used a common garden design to ensure the measurement of genetically based differences between individuals and focused on traits known to be heritable in sticklebacks (Bell 2005, Dingemanse *et al*. 2012, Karhunen *et al*. 2014). Predation elicited behaviours that could be considered to be advantageous in their corresponding environments and, particularly in the marine (ancestral) individuals. Indeed, in the presence of predators, fish would reduce activity time and foraging rates (thus decreasing their probability of mortality) while they increased these behaviours, and consequently their resource intake, in the absence of predators. Selection acting on this new advantageous variation in predator-free habitat would thus promote the evolution of bold behaviours. Nonetheless, Futuyma (2017) argued that “*phenotypic plasticity could be said to truly play a leading role (with genes as followers) if an advantageous phenotype were to be triggered by an environment that really is novel for the species lineage”*. In the case of *P. pungitius* – and more generally, in the case of predation – it is difficult to argue that the absence of predators is a truly novel condition to marine ancestors of freshwater adapted populations. Instead, the selection pressure imposed by predation in the wild could be viewed as a parameter with fluctuating intensity rather than a discrete state of the marine habitat (Moore *et al*. 2021). As such, varying levels of predation may have shaped the distribution of behaviours in ancestral populations of *P. pungitius* through balancing selection, and generated standing variation promoting local adaptation to freshwater habitats even through plastic responses. Although our results may not provide direct evidence for the role of plasticity in leading adaptive evolution, our study opens an interesting avenue of research to investigate the fitness effects of predation pressure in *P. pungitius*, and more generally, to consider the role of predation-induced plasticity in the evolution of complex traits.

There were marked behavioural differences between marine and pond sticklebacks and our findings are in agreement with those found in earlier studies (Herczeg *et al*. 2009a; Herczeg & Välimäki, 2011). However, in contrast to earlier studies (*e*.*g*., Herczeg et al. 2009a; Herczeg & Välimäki, 2011; Laine *et al*., 2014), all fish in our study were reared in groups. Since nine-spined sticklebacks display social behaviour such as schooling (Herczeg *et al*. 2009c), it is possible that the behaviours measured in our study were affected by this social component. Nonetheless, such social effects in the behaviours along the shy-bold continuum have been shown to exacerbate pre-existing differences in another fish species and was only found to affect shy individuals (*i*.*e*., shy individuals are shyer in the presence of shy conspecifics, Frost *et al*. 2007). Therefore, it is possible that shy behaviour (low exploration and risk-taking) was enforced in shy groups also in our study. This, however, might only accentuate existing behavioural differences, and would not have an effect on our conclusions. This is especially the case since the bold behaviour of the pond populations would have been relatively unaffected by group rearing. Overall, our large replicated common garden design provides robust evidence for the genetic basis of behavioural variation in wild stickleback populations from the two contrasting habitats.

Another important aspect of sociality in the expression of behaviours in *P. pungitius* is intraspecific competition. Indeed, the colonization of predator-free and low-productivity pond habitat is also associated with high levels of intraspecific competition and the evolution of gigantism and bold behaviours in the ponds has also been hypothesized to stem from this increased competition (Herczeg *et al*. 2009a,b). In such environments, the relaxation of predation pressure and absence of other species sharing similar trophic niche has inevitably led to the need for conspecifics to compete for limited food resources. Hence, it is possible that predation alone would not be sufficient to explain the evolution of bold behaviours and our current experimental setup does not allow to disentangle the effects of predation from the effects of intraspecific competition. However, an important result of our study is that the relaxation of predation pressure directly enhanced the foraging rate – a particularly important life-history trait – in all populations. Therefore, our results suggest that the relaxation of predation pressure would have allowed ‘quick and heavy’ feeders to acquire more resources in predator-free environments, which in turn, would be favoured by the new selection pressure imposed by the pond habitats. Future studies specifically testing for the interaction between predation risks and interspecific competition (*e*.*g*., Urban *et al*. 2015) are needed to shed more light on this specific aspect of behavioural evolution in *P. pungitius*.

In conclusion, we have demonstrated that genetically based differences in complex behaviour in Fennoscandian nine-spined sticklebacks have repeatedly evolved in similar environments and most likely in response to the same selection pressure. This provides strong evidence that this complex trait has evolved by natural selection in this species (cf. Schluter 2004). We also demonstrated that the phenotypically plastic response to the relaxation of predation pressure is aligned with the direction of evolutionary divergence observed in the wild, suggesting that phenotypic plasticity has likely contributed to the early stages of evolution of behaviour in freshwater habitats. Overall, our study shows that genetically determined behaviours can evolve through natural selection, and that behavioural traits are well suited to studying local adaptation in general.

## ETHICAL STATEMENT

All experiments were conducted under a permit from the Animal Experiment Board in Finland (permit reference ESAVI/4979/2018).

## AUTHOR CONTRIBUTION

AF and JM conceived the study; AF and EP conceived the experimental setup; EP performed all behavioural measurements; AF and EP analysed the data; AF led the writing of the manuscript with contributions from EP and JM; JM funded the study.

## DATA AVAILABILITY

Raw data will be deposited to Dryad upon acceptance. R code to reproduce the analyses will be made available at https://github.com/afraimout/.

## ACKNOWLEDGEMENTS

Thanks are due to Jacquelin DeFaveri, Niko Björkell, Karlina Ozolina, Niina Nurmi, and Miinastiina Issakainen for their precious help in sampling and rearing sticklebacks. We thank Xue-Yun Feng for his help in sampling perches and Nicolas Navarro for helpful comments on the phenotypic vector analysis. Our research was funded by the Academy of Finland (grant # 218343 to JM). The authors have no conflict of interest to declare.

