## supplementary material for "Relaxed risk of predation drives parallel evolution of stickleback behaviour"

**Table of contents**


| | |
|---|---|
| **Supplementary methods** | **Page 2-3** |
| **Figure S1. Schematic representation of the experimental aquaria.** | **Page 4** |
| **Figure S2. Kaplan-Meier survival analysis** | **Page 5** |
| **Table S1. Summary of samples** | **Page 6** |
| **Table S2. Results of the censored linear regression using the *censReg* R package.** | **Page 7** |
| **Table S3. Results of the linear mixed model with Box-Cox transformed data.** | **Page 8** |
| **Table S4. Pairwise comparisons of survival curves.** | **Page 9** |

### Supplementary methods

*Experimental aquaria*

Behavioural arenas were lined with polystyrene to prevent any visual disturbance from outside. Both aquaria had a clump of artificial algae (used as a refuge) and an opaque plastic cylinder with a small openable door in another corner against the same wall as the refuge (Fig. S1). The water in the control and predation treatment aquaria was not connected in any way so that fish in the control aquarium could not get any chemical nor visual cues from the perch. Apart from the absence of perch, the experimental aquarium of the control treatment was strictly identical to the one used in predation treatment. Within each tank, the water flowed from the holding arena to the behavioural arena, so that focal fish in the predation treatment could get chemical and visual cues from the perch. The perch were changed six times during the experimental period to prevent pseudo-replication as well as the habituation of the perch to the stickleback stimuli from having an effect on the results. A total of nine perch and seven unique perch pairs were used in the study. In the control treatment, the experimental aquarium housed no perch in the holding arena.

*Behavioural testing*

All behavioural testing were conducted over the course of 37 days in April-May 2019 divided into two temporal blocks, morning (8:30 – 12:30) and afternoon (13:00 – 18:00). At the time of testing, the mean age of the fish was 316.4 days with a standard deviation of 23.8 days. Since expression of exploration is known to differ between socially and solitarily reared fish (Jolles *et al*. 2016), tanks only housing a single fish were not assayed, so that all fish used in the experiments had been reared in a group. From each family, eight individuals were used for the experiments. If a family had less than eight fish, all individuals were used. Within each family, individuals were distributed evenly between treatments (predation and control, see above) and temporal blocks (morning and afternoon), so that each group had representation from both of the replicates in a family. If the family had less than 8 individuals, individuals were assigned into groups to maximize variation between (i) treatment, (ii) tank, and (iii) temporal block. All individuals were distributed in a random order across the experimental days. Both exploration behaviour variables (time-to-head-out and time-to-body-out) were highly correlated ($\rho$ = 0.951 [0.941; 0.959], $p < 0.001$) and the arithmetic mean of both variables was used as exploration time in our analyses.

*Data analysis of right-censored data*

We used 3 different statistical approaches to model our right-censored, time-to-event variables. First, we used the Kaplan-Meier survival analysis framework (Crowley & Breslow 1984) and fitted survival curves using the *survival* (v.3.2-11, Therneau 2021) and *survminer* R packages (v.0.4.9, Kassambara *et al.* 2017). Because the event measured in our time-to-event data is the expression of a behaviour rather than actual death or survival as is usually the case in such analyses, the survival curves we estimated correspond to the expected proportion of fish having expressed the behaviour of interest at a certain time. Original time-to-event variables were binary transformed by assigning a value of 0 to all individuals with the maximum value of 300s and 1 to all other individuals. We used the *survfit* function and included habitat of origin and treatment as categorical variables to estimate the survival curve for each behaviour.

We then compared the differences in behaviour between habitats and treatments by computing the differences between survival curves using the log-rank test (Kleinbaum & Klein, 2012) implemented in the *pairwise_survdiff* function of the *survminer* package and using the Bonferroni *p*-value adjustment method.

Second, we applied a Box-Cox transformation (Box & Cox 1964) to all right-censored measurements and using the *boxcox* function of the *MASS* R package (v.7.3-54, Venables & Ripley 2002) used the transformed data as response variables in a Linear Mixed Model framework. We used the *lmer* function of the *lme4* R package (v.1.1-27, Bates *et al.* 2015) and fitted a model with habitat and treatment and their interaction as fixed effects. Temporal block and age-corrected body size were used as covariate and the perch pair and tank identities were used as random effects. We tested for the significance of the fixed effects using Wald chi-square tests using the *Anova* function in the *car* R package (v.3.0-11, Fox & Weisberg 2019).

Finally, we used the *censReg* R package to fit censored regression to our data, as described in the main text.

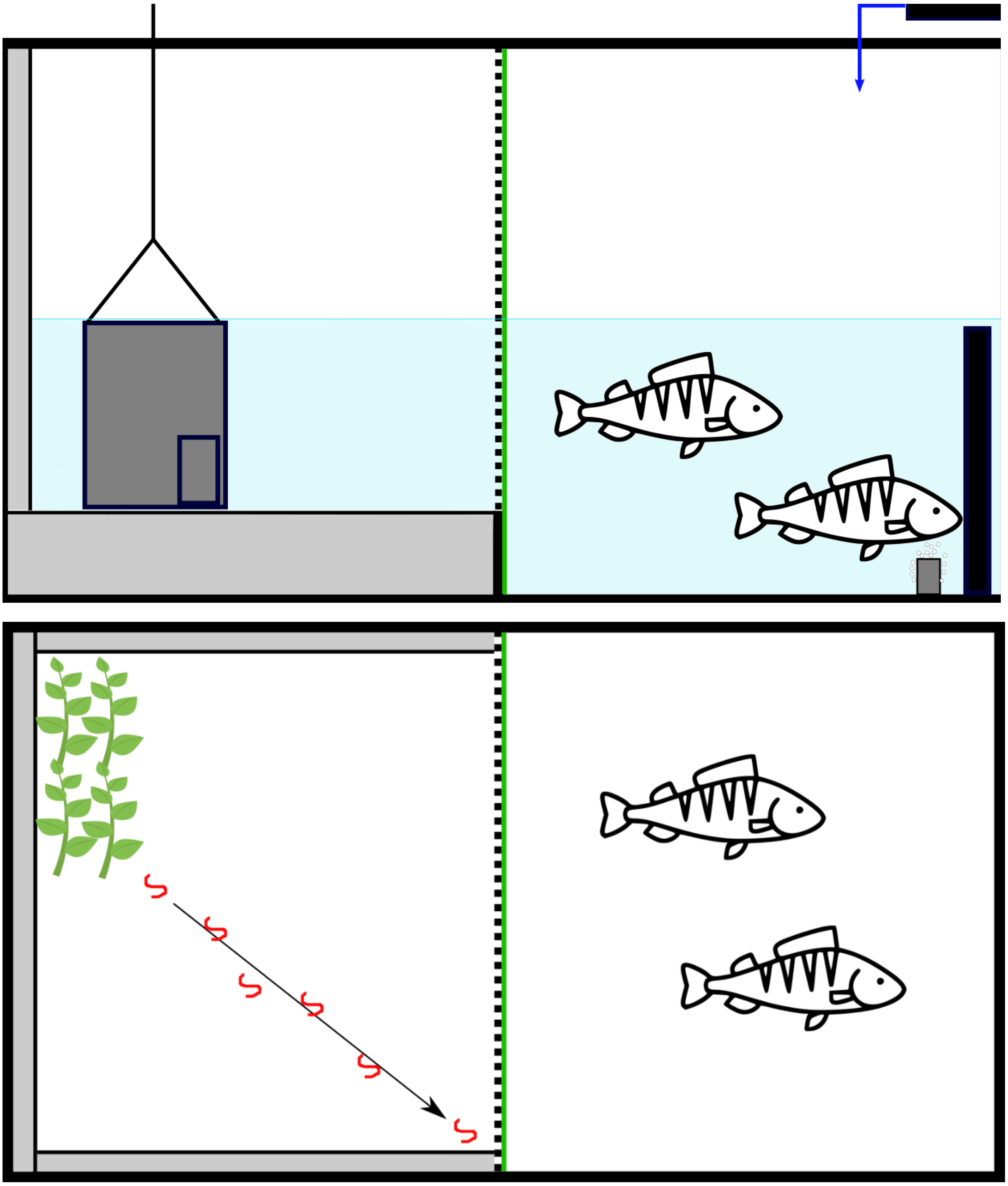

**Fig. S1. Schematic representation of the experimental aquaria.** Upper panel shows a lateral view of the experimental aquarium during the exploration test. The left side corresponds to the behavioural arena including the polystyrene lining (grey rectangles) and the opaque plastic cylinder used for exploration trials. The right side corresponds to the holding arena housing the pair of perch (for predation treatment), the water inlet (blue arrow) and outlet (black rectangle) as well as an air stone (grey rectangle). Lower panel shows a top view of the experimental aquarium in the risk-taking test. A clump of artificial algae was set up as a refuge in the behavioural arena (top left corner). Chironomid larvae (food item, in red) were pipetted into the tank following a diagonal line (black arrow). A transparent non-hermetic plastic divider (transversal green and dashed black lines) was set up to allow visual and chemical cues between behavioural and holding arenas. Perch were introduced in the holding arena (right side) in the predation treatment only.

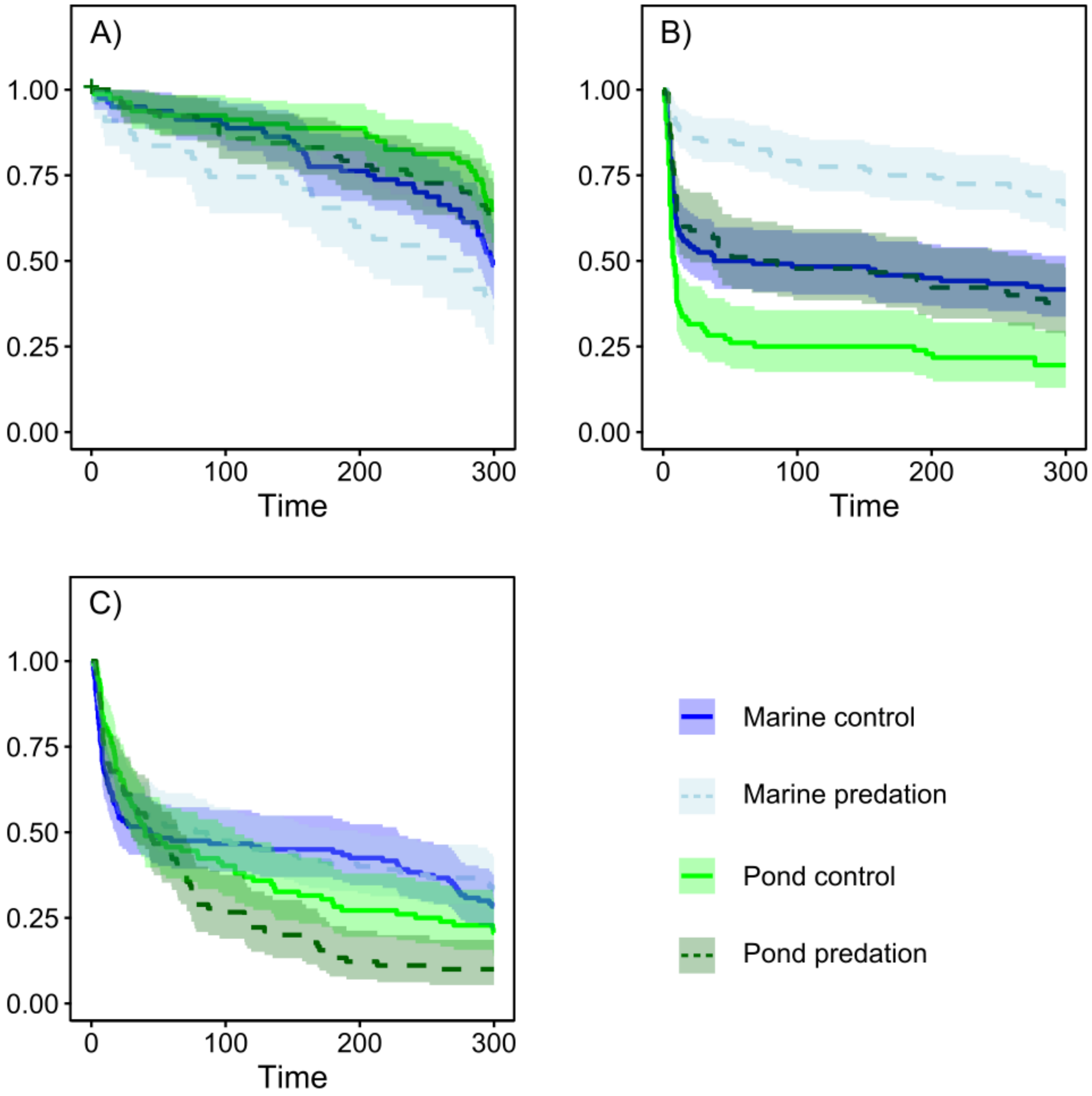


**Fig. S2. Kaplan-Meier survival analysis of time-to-event variables.** Survival curves for the control treatment (solid lines) and predation treatment (dashed lines) are shown with their 95% confidence interval (shading) for marine (blue) and pond (green) individuals. The x-axis represents the total time of measurement of the experiment. The y-axis corresponds to the proportion of individuals active in the open area (A); the proportion of fish staying in the refuge after food administration (B) and the proportion of fish staying in the refuge during the exploration trial (C).

**Table S1. Summary of samples.**

| Code | Habitat | Country | Sampliong location | GPS coordinates | Number of families | Total number of individuals |
|---|---|---|---|---|---|---|
| FIN-TVA | Marine | Finland | Tvärminne | 59.83333, 23.2 | 10 | 75 |
| FIN-POR | | | Pori | 61.59111, 21.47295 | 10 | 71 |
| FIN-RAA | | | Raahe | 64.68818, 24.46189 | 5 | 40 |
| SWE-UME | | Sweden | Umeå | 63.63739, 19.99438 | 10 | 74 |
| FIN-PYÖ | Freshwater | Finland | Pyöreälampi | 66.26226, 29.42916 | 5 | 49 |
| FIN-RYT | | | Rytilampi | 66.38482, 29.31561 | 10 | 76 |
| FIN-KRK | | | Kirkasvetinenlampi | 66.43673, 29.13568 | 5 | 30 |
| SWE-BYN | | Sweden | Bynästjärnen | 64.45416, 19.44075 | 10 | 46 |

**Table S2. Results of the linear regressions using the original data.** Trait type**:** the two categories of behaviours. Trait: the behavioural trait used as response variable. Method: the statistical framework used to model the data (see *Material & Methods* in the main text). Effect: name of the fixed effect. Estimate: coefficient of the fixed effect. Std. Err.: standard error of the coefficient. T: *t*-value. P: *p*-value. Bold values represent significant differences between curves ($p$ < 0.05) and italic values correspond to non-significant differences. Colons indicate fixed terms interaction.

| **Trait type** | **Trait** | **Method** | **Effect** | **Estimate** | **Std. Err.** | **T** | **P** |
|---|---|---|---|---|---|---|---|
| Exploration | Emergence time | censReg | Intercept | 166.761 | 16.989 | 9.815 | **<2e-16** |
| | | | Size | 3.421 | 1.768 | 1.934 | *0.053* |
| | | | Habitat | 47.324 | 23.769 | -1.991 | **0.046** |
| | | | Treatment | 7.870 | 20.705 | 0.380 | *0.704* |
| | | | Block | -0.281 | 15.496 | -0.018 | *0.985* |
| | | | Habitat:Treatment | -47.197 | 31.230 | -1.511 | *0.131* |
| Foraging | Open time | censReg | Intercept | 169.535 | 39.064 | 4.340 | **1.43e-05** |
| | | | Size | -3.857 | 4.203 | -0.918 | *0.359* |
| | | | Habitat | 230.835 | 58.107 | 3.973 | **7.11e-05** |
| | | | Treatment | -188.547 | 49.161 | -3.835 | **1.25e-04** |
| | | | Block | -26.929 | 36.674 | -0.734 | *0.463* |
| | | | Habitat:Treatment | 144.684 | 74.714 | 1.937 | *0.053* |
| | Risk taking | censReg | Intercept | 216.096 | 23.115 | 9.349 | **< 2e-16** |
| | | | Size | 8.15319 | 2.435 | 3.348 | **8.15e-04** |
| | | | Habitat | -140.009 | 31.442 | -4.453 | **8.47e-06** |
| | | | Treatment | 156.757 | 29.373 | 5.337 | **9.46e-08** |
| | | | Block | -2.591 | 21.102 | -0.123 | *0.902* |
| | | | Habitat:Treatment | -57.342 | 42.472 | -1.350 | *0.177* |
| | Feeding | GLM | Intercept | 2.131 | 0.179 | 11.922 | **< 2e-16** |
| | | | Size | -0.028 | 0.019 | -1.496 | *0.135* |
| | | | Habitat | 0.585 | 0.251 | 2.326 | **0.020** |
| | | | Treatment | -1.293 | 0.222 | -5.804 | **6.49e-09** |
| | | | Block | -0.059 | 0.166 | -0.356 | *0.722* |
| | | | Habitat:Treatment | 0.534 | 0.335 | 1.593 | *0.111* |

**Table S3. Results of the linear mixed model with the Box-Cox transformed time-to-event data.** Results from the Wald Chi-square ($\chi^2$) test of fixed effects significance are. Trait: the behavioural trait used as response variable. Effect: name of the fixed effect. LRT: Likelihood-Ratio Test. P: *p*-value. Bold values represent significant differences between curves ($p < 0.05$) and italic values correspond to non-significant differences. Colons indicate fixed terms interaction.

| Trait | Effect | LRT ($\chi^2$) | P |
|---|---|---|---|
| Emergence time | Size | 4.9300 | **0.026** |
| | Habitat | 3.3392 | *0.067* |
| | Treatment | 0.0015 | *0.968* |
| | Block | 0.0148 | *0.903* |
| | Habitat:Treatment | 2.3042 | *0.129* |
| Open time | Size | 0.5366 | *0.463* |
| | Habitat | 54.3679 | **1.663e-13** |
| | Treatment | 12.9993 | **3.116e-04** |
| | Block | 0.0930 | *0.760* |
| | Habitat:Treatment | 6.0426 | **0.014** |
| Risk taking | Size | 14.5395 | **1.372e-04** |
| | Habitat | 55.0803 | **1.157e-13** |
| | Treatment | 55.3359 | **1.016e-13** |
| | Block | 0.0057 | *0.939* |
| | Habitat:Treatment | 0.1372 | *0.711* |

**Table S4. Pairwise comparisons of survival curves.** Survival curves for each behaviour and each habitat x treatment pair were compared using a pairwise log-rang test. Bold values represent significant differences between curves ($p < 0.05$) and italic values correspond to non-significant differences.

| **Trait** | | Marine control | Marine predation | Pond control |
|---|---|---|---|---|
| Emergence time | Marine predation | *0.421* | **-** | **-** |
| | Pond control | *0.199* | **0.001** | **-** |
| | Pond predation | *0.940* | **0.015** | *1.000* |
| Open time | Marine predation | *1.000* | **-** | **-** |
| | Pond control | *1.000* | *0.9496* | **-** |
| | Pond predation | **0.045** | **0.005** | *0.315* |
| Risk taking | Marine predation | **3.5e-05** | **-** | **-** |
| | Pond control | **0.001** | **2.5e-16** | **-** |
| | Pond predation | *1.000* | **8.6e-06** | **0.002** |
